# Phosphatidic acid inhibits SNARE priming by inducing conformational changes in Sec18 protomers

**DOI:** 10.1101/365775

**Authors:** Matthew L. Starr, Robert P. Sparks, Logan R. Hurst, Zhiyu Zhao, Andres Arango, Muyun Lihan, Jermaine L. Jenkins, Emad Tajkhorshid, Rutilio A. Fratti

**Author notes:** These authors contributed equally (Matthew L. Starr, Robert P. Sparks).

## Abstract

Eukaryotic homeostasis relies on membrane fusion catalyzed by SNARE proteins. Inactive SNARE bundles are re-activated by Sec18/NSF driven disassembly to enable a new round of fusion. We previously found that phosphatidic acid (PA) binds Sec18 to sequester it from SNAREs. Dephosphorylation of PA dissociates Sec18 from the membrane allowing it to engage SNARE complexes. We now report that PA induces conformational changes in Sec18 protomers, while hexameric Sec18 cannot bind PA membranes. The association of Sec18 with PA was shown to be sensitive to membrane curvature, suggesting that regulation could vary on different organelles in a curvature dependent manner. Molecular dynamics showed that PA binding sites exist on the D1 and D2 domains of Sec18 and that residues needed for binding were masked in the hexameric form of the protein. Together these data indicate that PA regulates Sec18 function through altering protein architecture and stabilizing membrane-bound protomers.

## INTRODUCTION

Membrane fusion is necessary for all eukaryotes to effectively transport cellular components between organelles. Vesicle trafficking is carried out through a series of events that are highly conserved across eukarya (Jahn and Sudhof, 1999). Many proteins that drive the process differ between eukaryotic species, but all perform similar roles allowing compartment contact, bilayer fusion, and luminal content mixing (Jahn et al., 2003). The final stage of membrane fusion, and luminal content mixing, is catalyzed by SNARE proteins. Each participating membrane contributes either an R-SNARE or three Q-SNARE coils that wrap around each other to form a parallel four-helical *trans-SNARE* complex that brings membranes into close apposition. The formation of such complexes releases free energy that is transmitted to the membranes to trigger fusion. Once fusion occurs and membranes are merged, the four helical SNARE bundle, now a cis-SNARE complex, is inactive and requires reactivation in order to undergo a new round of fusion.

The activation cis-SNAREs, also known as *Priming*, is carried out by the AAA+ protein Sec18/NSF and its adaptor protein Sec17/α-SNAP (Mayer et al., 1996). Current models suggest that NSF primes *cis*-SNAREs through a “loaded-spring” mechanism triggered by cis-SNARE recognition and ATP hydrolysis (Ryu et al., 2015). NSF binds to cis-SNAREs with the help of α-SNAP to form what is known as the 20S complex (Chang et al., 2012; Sollner et al., 1993; Wilson et al., 1992; Zhao et al., 2015). In its active form, NSF forms a homohexamer which surrounds the cis-SNAREs and α-SNAP proteins to form the 20S particle (Fleming et al., 1998). Association with cis-SNARE-α-SNAP complexes triggers ATP hydrolysis which leads to a large conformational change in the protein. This generates enough force to disrupt the 20S complex and separate the individual SNAREs from each other effectively reactivating them.

Previous work identified that both NSF and Sec18 bind to the regulatory glycerophospholipid phosphatidic acid (PA) (Manifava et al., 2001; Starr et al., 2016). PA has been shown to have regulatory effects in multiple vesicular trafficking pathways including sporulation, regulated exocytosis, lysosomal maturation, and homotypic vacuole fusion (Liu et al., 2007; Nakanishi et al., 2006; Rogasevskaia and Coorssen, 2015; Sasser et al., 2012; Starr et al., 2016). In the case of Sec18, increased PA levels lead to reduced priming activity likely due to a decrease in recruitment to *cis*-SNAREs (Starr et al., 2016). On yeast vacuoles, PA is converted to diacylglycerol (DAG) by the PA phosphatase Pah1, an ortholog of mammalian Lipin1. In the absence of Pah1 activity, PA levels remain intact and sequester Sec18 from *cis*-SNARE complexes to prevent priming and arrest the fusion pathway (Sasser et al., 2012). DAG can be converted to PA through the action of the DAG kinase Dgk1, whose inactivation leads to elevated DAG concentrations that enhance fusion through modulating the activity of the Rab GTPase Ypt7 (Miner et al., 2017). Thus, the interconversion of PA and DAG serves as a regulatory switch to control vacuole fusion.

Here we asked what effects PA-binding has on the overall architectural dynamics of Sec18/NSF that could lead to a decrease in its priming activity. To do so, we measured binding of monomeric and hexameric Sec18 to different forms of PA. We report that monomeric Sec18 has significantly stronger binding than the hexameric form to all forms of PA. We probed changes to the architecture of Sec18 when bound to short-chain PA and found that the protein exists in a significantly different conformation in its PA-bound state, without significant changes to its secondary structure. To study the mechanism of Sec18 binding to PA, molecular dynamics simulations were performed using the mammalian version of Sec18, namely NSF. NSF was used as it has high identity to Sec18 and has more structural information available at the protein data bank (PDB ID: 3J94) (Zhao and Brunger, 2016). The molecular dynamic simulations performed suggest NSF binds to PA at regions of the protein that are only exposed in the monomeric state of the protein. Taken together, we propose that PA regulates the priming activity of NSF/Sec18 by limiting the formation of its active hexamer.

## RESULTS

### Sec18 monomer binds to PA with higher affinity than the hexameric form

Our previous work showed that Sec18 preferentially bound to liposomes containing phosphatidylcholine (PC), phosphatidylethanolamine (PE) and PA relative to those composed of only PC and PE, or ones where PA was replaced with DAG (Starr et al., 2016). This was in keeping with older findings showing that mammalian NSF bound to resin-linked PA (Manifava et al., 2001). Here our studies were extended to further define how Sec18 binds to PA. To start we used microscale thermophoresis (MST) to acquire binding affinities to dioctonyl PA (C8-PA), which prevents Sec18 from binding cis-SNARE complexes, consequently precluding priming from occurring (Starr et al., 2016). We used both monomeric and hexameric Sec18 with a range of C8-PA. The C-terminal 8xHistidine tag of Sec18 was labeled with Ni-NTA Atto 488. As shown in Figure 1A, monomeric Sec18 (mSec18) bound to C8-PA with a *K*_*D*_ of 1.4±0.68 μM (blue circles), whereas the hexameric form (hSec18) had a *K*_*D*_ of 29±8.6 μM (red squares). This suggested that either hSec18 has residues occluded for PA binding or is in an inappropriate conformation to bind C8-PA in hexameric form. It is possible that a small soluble C8-PA could access a binding site on Sec18 that is obscured in the hexamer, where membrane contained PA is unable to reach PA binding regions on Sec18 when Sec18 is hexamerized, especially regions contained in the hexamerization interface of Sec18 hexamer.

**Figure 1.**
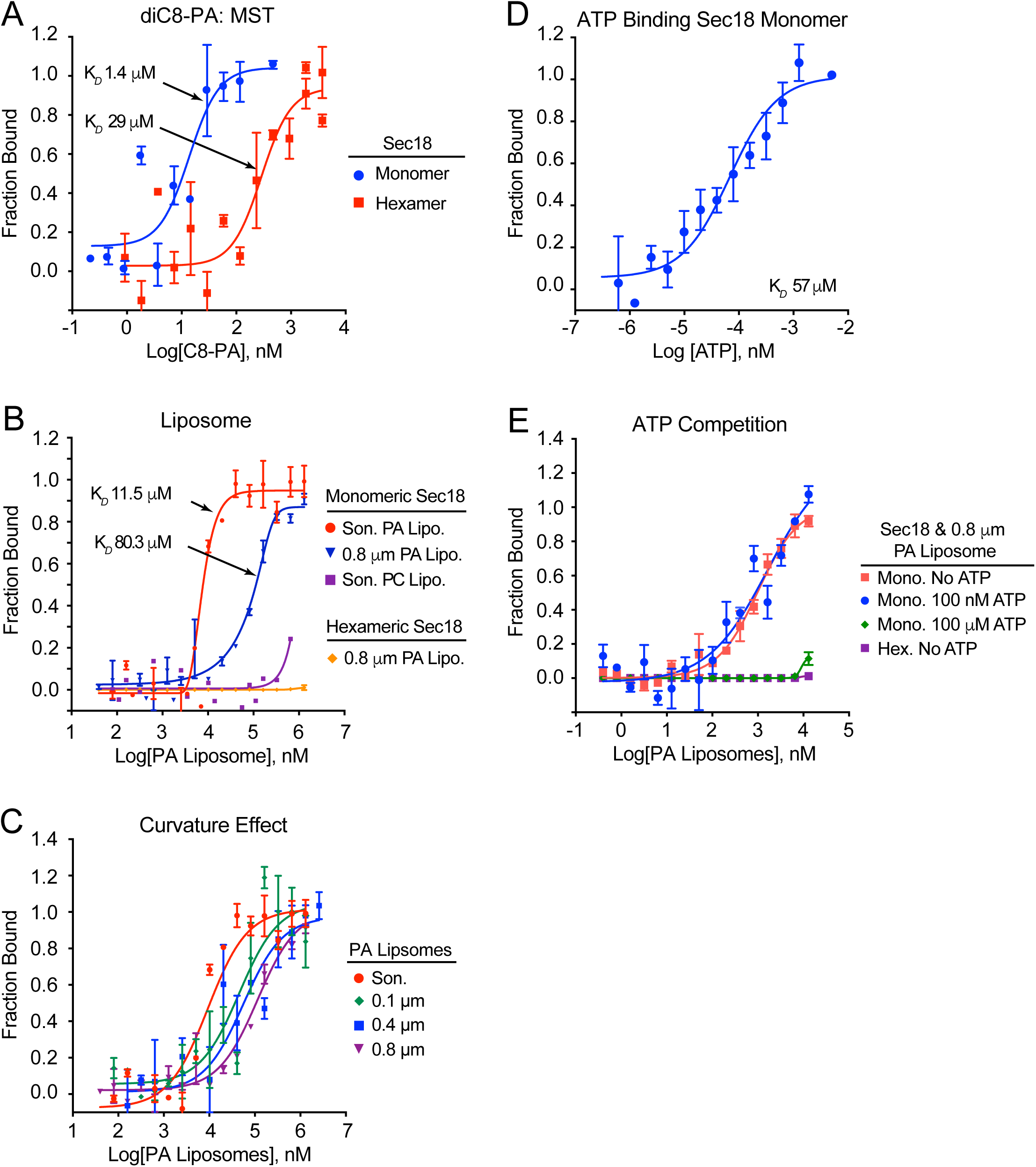
Sec18 Hexamer and monomer binding affinity for PA. (**A**) C8-PA MST measurements were performed using purified Sec18 monomer and hexamer labeled with Ni-NTA Atto-488 dye in the blue channel at 90% LED and High MST using NT.115 Labeled Thermophoresis. Binding affinity was measured using thermophoresis at 15 sec mixing separate reactions of half 100 nM Atto 488 labeled Sec18 monomer and half 1:1 titrations of C8-PA with highest concentration 370 μM according to Graphpad Sigmoidal 4PL curve. (**B**) The *K*_*D*_ of ATP for Sec18 monomer was measured using labeled Sec18 monomer with Ni-NTA Atto 488 dye as in Fig. 1A with 1:1 titrations of ATP Solution in PBS. (**C**) The *K*_*D*_ of Sec18 monomer and hexamer to sonicated and 800 nm diameter PA liposomes (10% PA, 70% PC, and 20% PE) and PC liposomes (80% PC, 20% PE) was measured using MST as in Fig. 1A. Concentrations of lipid of 1.3 mM maximum were titrated as in Fig. 1A for both PA and PC liposomes and affinities for both Sec18 hexamer binding to PA liposomes as well as monomer binding to PC liposomes was not measurable as they did not saturate. (**D**) ATP competition with mSec18 binding to 800 nm PA liposomes was measured as in Fig. 1A was measured using MST ATP concentrations of 100 nM (D2 saturating) and 100 μM (D1 saturating) and compared to the *K*_*D*_ of monomer and hexamer in the absence of ATP. (**E**) Sec18 monomer affinity for different size of PA liposomes was measured using MST as in Fig. 1A for sonicated, 100 nm, 400 nm and 800 nm. Curve fitting was performed using Graphad sigmoidal 4PL curve All measurements taken at 15 s thermophoresis using M.O. Affinity Analysis software as in Fig. 1A.

Due to the difference in binding affinities to C8-PA, we next asked if limiting the mobility of PA to two dimensions would show a similar disparity between the monomer and hexamer. To this aim we used sonicated liposomes as previously reported as well as extruded 0.8 μm diameter liposomes to approximate the diameter of yeasts vacuoles. We found that mSec18 bound sonicated PA liposomes with a *K*_*D*_ of 9.3±1.32 μM (red circles) and 0.8 μm PA liposomes with a *K*_*D*_ of 97.7±10.9 μM (blue triangles) (Fig. 1B). In both cases, mSec18 bound with a lower affinity relative to C8-PA, supporting the notion that membranous PA is limited in its interactions with Sec18. As a control for PA specificity we tested sonicated liposomes containing only PC and PE (purple squares), and found that Sec18 did not appreciably bind, which is in keeping with our previous study. Importantly, we found that hSec18 did not bind 0.8 μm PA liposomes (orange diamonds) or 0.1 μM PA liposomes (not shown). This data is indicative of two major conclusions. First, it is apparent that membrane curvature affects Sec18 binding to PA, similar to other PA binders (Putta et al., 2016). Second, and more importantly, is that the hexameric Sec18 lacks the ability to bind PA, potentially by masking a binding site or by restricting conformational changes needed to bind PA.

### Membrane curvature affects Sec18 binding to PA

To further test the role of membrane curvature we generated PA liposomes using extrusion with diameters of 0.1, 0.4 and 0.8 μm. These were used in parallel to sonicated liposomes with an average diameter of 30-50 nm (Lapinski et al., 2007). MST experiments showed that the affinity for PA was reduced as the diameter of membranes increased. Sonicated PA liposomes bound mSec18 with a *K*_*D*_ of 9.3±1.32 μM, while the *K*_*D*_ values of 0.1 μm, 0.4 μm and 0.8 μm liposomes were 45.1±11.8 μM, 56.8±15.6 μM, and 97.7±10.9 μM, respectively (Fig. 1C). Although not a linear effect, it is clear that membrane curvature alters the ability of Sec18 to bind PA. This also suggests that the effect of PA on Sec18 mediated SNARE priming could vary depending on the local curvature of an organelle.

### ATP can reduce PA binding by Sec18

Sec18/NSF like many other AAA+ proteins contains two nucleotide binding domains (NBD) each residing in a one of the domains that make up the rings of the hexameric protein. The D1 ring of Sec18 hydrolyses ATP to generate the mechanical force needed to disrupt cis-SNARE bundles whereas the D2 ring binds ATP to stabilize the hexameric form of the protein. This is reflected in the different affinities for ATP found between the two NBDs. In NSF the D1 NBD binds ATP with a *K*_*D*_ of 15-20 μM, while the D2 NBD binds with a *K*_*D*_ of 30-40 nM (Matveeva et al., 1997). Here we asked ATP binding, which is linked to large conformational changes during SNARE priming, would affect PA binding. First we determined the affinity of mSec18 for ATP using MST and found that mSec18 had a *K*_*D*_ of 56±16 μM, which likely reflects the low-affinity binding site in D1 (Fig. 1D). We were unable to detect the high affinity binding to be expected of the D2 NBD. We then tested if ATP binding altered PA binding. mSec18 was pre-incubated with 0.8 μm PA liposomes, then introduced to 100 nM ATP, 100 μM ATP, or buffer alone (no ATP). This showed that in the presence of 100 nM ATP, a saturating concentration for the D2 NBD (not shown), there was no effect on PA binding as the curve overlapped with the no ATP control (Fig. 1E, blue circles vs red squares, respectively). Both conditions bound to PA with a *K*_*D*_ of approximately 100 μM. In contrast, saturating both NBDs with 100 μM ATP completely blocked binding to PA liposomes (green diamonds). As a negative control we incubated hSec18 with PA liposomes in the absence of ATP, which showed no binding at concentrations of PA liposomes tested (purple squares).

### Sec18 binds PA with similar affinity to DEP PA binding domain

In our previous study we competed Sec18 binding to PA liposomes with GST-DEP, a well characterized PA binding domain from the murine protein Dvl2 (Capelluto et al., 2014). Here we compared the binding affinity of DEP to mSec18. Because we previously used GST-DEP, we measured its binding in comparison to GST-Sec18. We also wanted to verify our MST data with surface plasmon resonance (SPR) and PA nanodiscs (PA-ND), which were linked to Ni-NTA SPR chips through the 6xHis tags of the ND scaffold proteins. These experiments showed that GST-Sec18 bound to PA-ND with a *K*_*D*_ of 2.7±2 μM (Fig. 2A), whereas GST-DEP had a *K*_*D*_ of 18±2 μM (Fig. 2B), indicating that Sec18 binds PA with a higher affinity relative to the *bona fide* PA-binding domain DEP. This was also seen using MST, where GST-Sec18 bound to PA-ND with a *K*_*D*_ of 0.6±.096 μM whereas GST-DEP had a higher *K*_*D*_ of 2.4±0.42 μM, and N Domain had a *K*_*d*_ of 8.5±14.4 μM. The increased affinity for PA nanodiscs is possibly due to the fact that nanodiscs as opposed to liposomes are quantified by actual number of nanodiscs and not total lipid concentration as in C8-PA and liposome measurements respectively. Alternatively, increased affinity of PA binding in comparison to the values in Figure 1 are attributed to the effect of GST dimerization or GST stabilization of conformation, which could contribute to an increase in avidity relative to His-tagged mSec18, similar what was seen with the protease inhibitor cystatin (Tudyka and Skerra, 1997).

**Figure 2.**
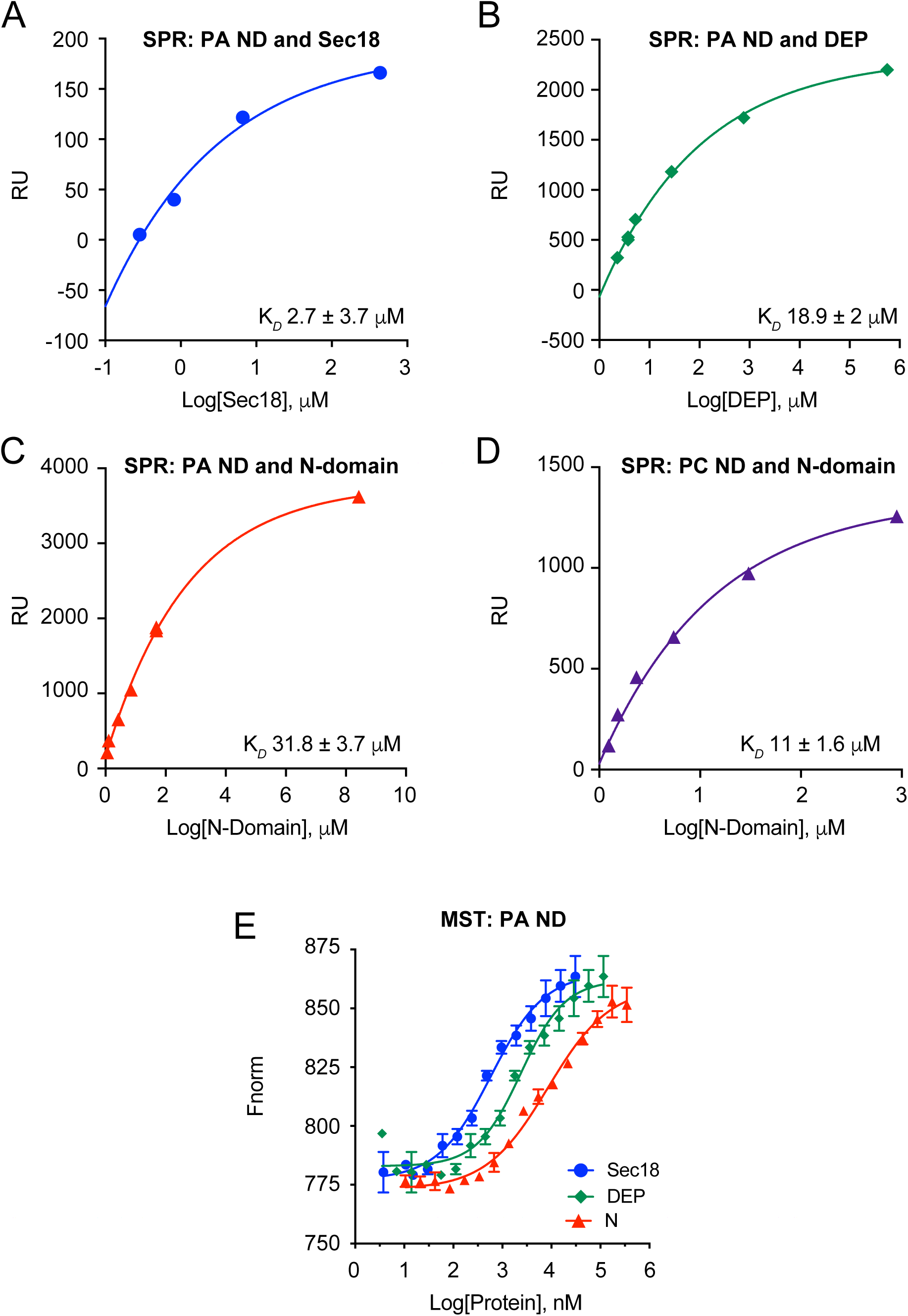
Sec18 Binding Affinity Compared to DEP PA Binding Domain for PA Nanodiscs. (**A**) SPR analysis of Sec18 monomer was performed with approximately 2000 RU of 5% PA nanodiscs attached to a Ni-NTA chip using a Biacore 300 Ni-NTA with flowrate 20 μL/s. The steady state fit was exported from BIAevaluate software to GraphPad at 4 seconds before injection stop set at 90 s with disassociation of 120s. (**B**) SPR Analysis of DEP with 5% PA nanodiscs. (**C**) SPR analysis of N domain for Sec18 monomer with 5% PA nanodiscs. (**D**) SPR analysis of PC nanodiscs for Sec18 monomer. (**E**) MST performed with mSec18, DEP, and N-domain with 100 nM Ni-NTA Atto 488 labeled 5% PA nanodiscs using 90% LED and 60% MST. M.O. Affinity analysis software was used and thermophoresis exported at 15 s.

Due to the PA inhibition of Sec18 binding to cis-SNARE complexes, we asked if the Sec18 N-terminal domain could bind to PA by itself. This notion is supported by the crystal structure of the NSF N-terminal domain showing the presence of a positive polybasic surface adjacent-to and lining the α-SNAP binding groove (Yu et al., 1999). SPR measurements of GST-N-domain from Sec18 binding to PA-ND showed a *K*_*D*_ of 31.8±3.7 μM **(Fig. 2C)**. Interestingly, the N-domain bound ND containing only PC and PE with a *K*_*D*_ of 11±1.6 μM **(Fig. 2D)**, suggesting that the N-domain has no lipid-binding specificity. Because full length Sec18 binds to PA and not PC, we conclude that the N-domain does not contribute to the regulatory association with PA. This is further supported by our MST data showing that mSec18 binds to PA-ND with an affinity that is nearly two orders of magnitude higher relative to the N-domain alone (Fig. 2E, blue circles vs red triangles).

### Phosphatidic acid alters the conformation of Sec18

Our data thus far suggests that Sec18 undergoes conformational changes that allow mSec18 to bind PA while hSec18 lacks the ability to bind the lipid. To further probe for conformational changes to Sec18 we tested whether PA significantly alters binding of 8-Anilino-1-naphthalenesulfonic acid (ANS) to Sec18. ANS is a dye that has been extensively used to test lipid-binding proteins because it associates with solution exposed hydrophobic motifs (Heyduk and Lee, 1989; Roberts et al., 1999). Binding of ANS to a protein results in an increase in fluorescence yield and a blue-shifted emission. Because we have previously seen PA binding to mSec18 we expected ANS to also bind the protein in our assay. As expected, we observed ANS binding to mSec18 in a dose-dependent fashion **(Fig. 3A-B)**. We next wanted to test for any conformational changes upon PA binding that altered ANS binding to Sec18. To do this, we titrated increasing amounts of C8-PA into our assay and measured changes in the ANS fluorescence spectra. Because C8-PA is partially hydrophobic, ANS was first incubated with each lipid concentration to obtain a background spectrum before protein was then added to the assay, and fluorescence was again measured. The difference spectra from these measurements shows that addition of C8-PA increases the binding of ANS to Sec18 **(Fig. 3C and F)**. To verify that the changes in ANS fluorescence were specific to PA binding, we tested the addition of DAG, the product of Pah1 activity on PA. No change in ANS fluorescence was detected in the presence of C8-DAG, which is consistent with inability of Sec18 to bind to DAG **(Fig. 3D and F)**. We also tested the anionic lipid phosphatidylserine (PS). Similar to what we observed with DAG, the addition of C8-PS had no effect on ANS fluorescence **(Fig. 3E and F)**. *In toto* these data suggest that C8-PA binding to Sec18 results in a conformational change in the protein that exposes additional hydrophobic pockets to solution. Such a change may account for the differences previously seen in Sec18 priming activity and cis-SNARE association (Starr et al., 2016).

**Figure 3.**
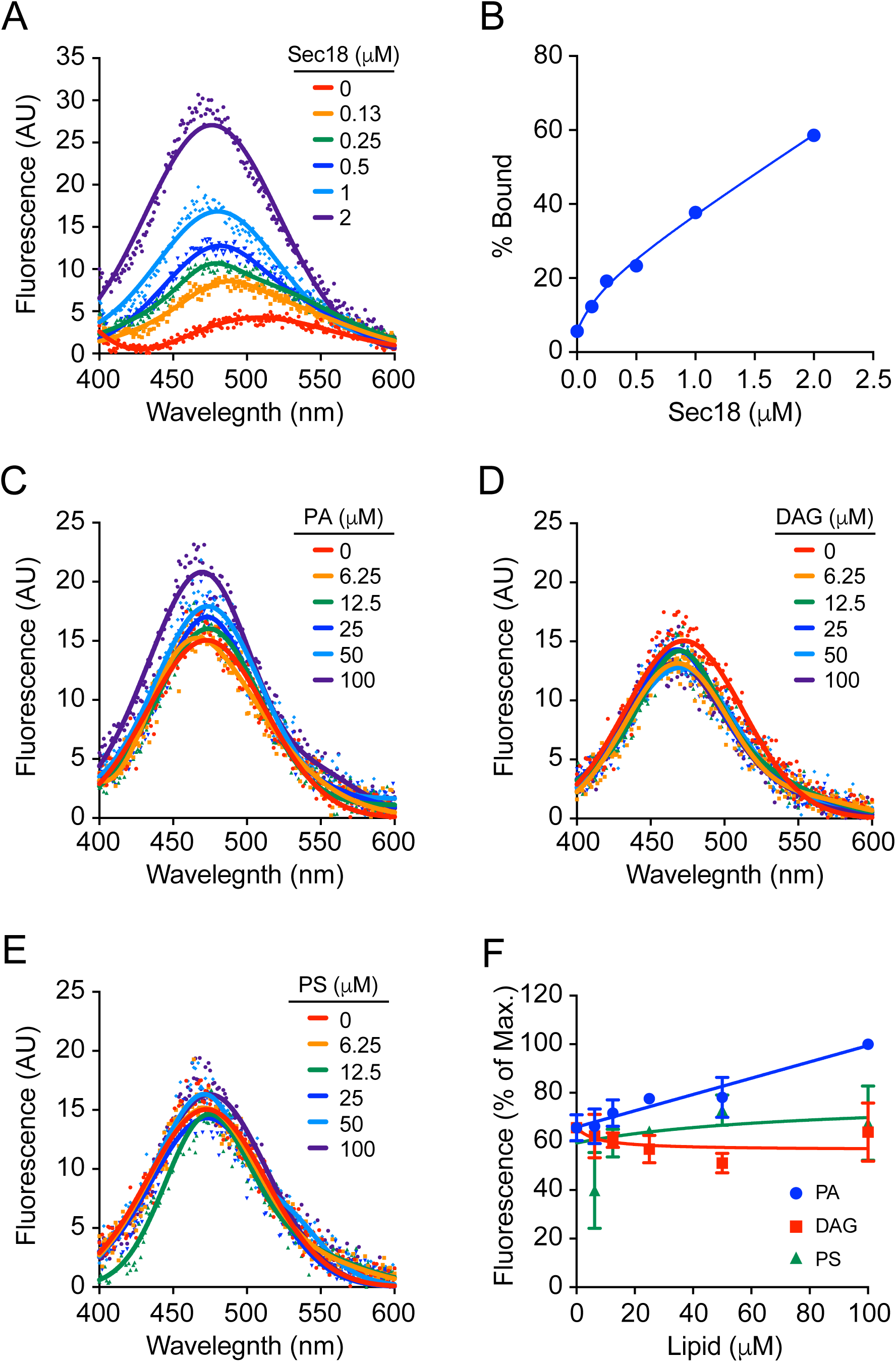
Short chain phosphatidic acid alters the binding of 1,8 ANS to Sec18. Increasing concentrations of Sec18_His8_ were incubated with ANS (5 μM) in assay buffer and a representative fluorescence spectrum (ex. 390, em. 400-600 nm) is shown (**A**). Relative fluorescence at 460 nm (**B**). Sec18_His8_ (0.5 μM) was incubated with increasing concentrations of short-chain lipids in the presence of ANS (5 μM) and fluorescence spectra were taken (ex. 390, em. 400-600 nm). A representative spectrum for each lipid tested is shown: C8-PA (**C**), C8-DAG (**D**), and C8-PS (**E**). (**F**) Maximum fluorescence for each lipid concentration was normalized against overall maximum fluorescence (100 μM C8-PA) for relative comparison.

To further probe for conformational changes to Sec18 induced by PA we utilized a limited proteolysis assay. Proteins can exhibit differences in their proteolytic cleavage profiles when bound to a ligand that significantly changes their overall architecture (Heyduk and Lee, 1989). Because we observed an increase in solution exposed regions of Sec18 in the presence of PA, *i.e*. increased ANS fluorescence, we expected to also see an increased sensitivity to protease degradation in the same conditions. To measure this, mSec18 was incubated with increasing concentrations of trypsin with and without C8-PA addition. As expected, mSec18 sensitivity to trypsin degradation increased in the presence of C8-PA, whereas the presence of DAG had no effect **(Fig. 4A-B)**.

**Figure 4.**
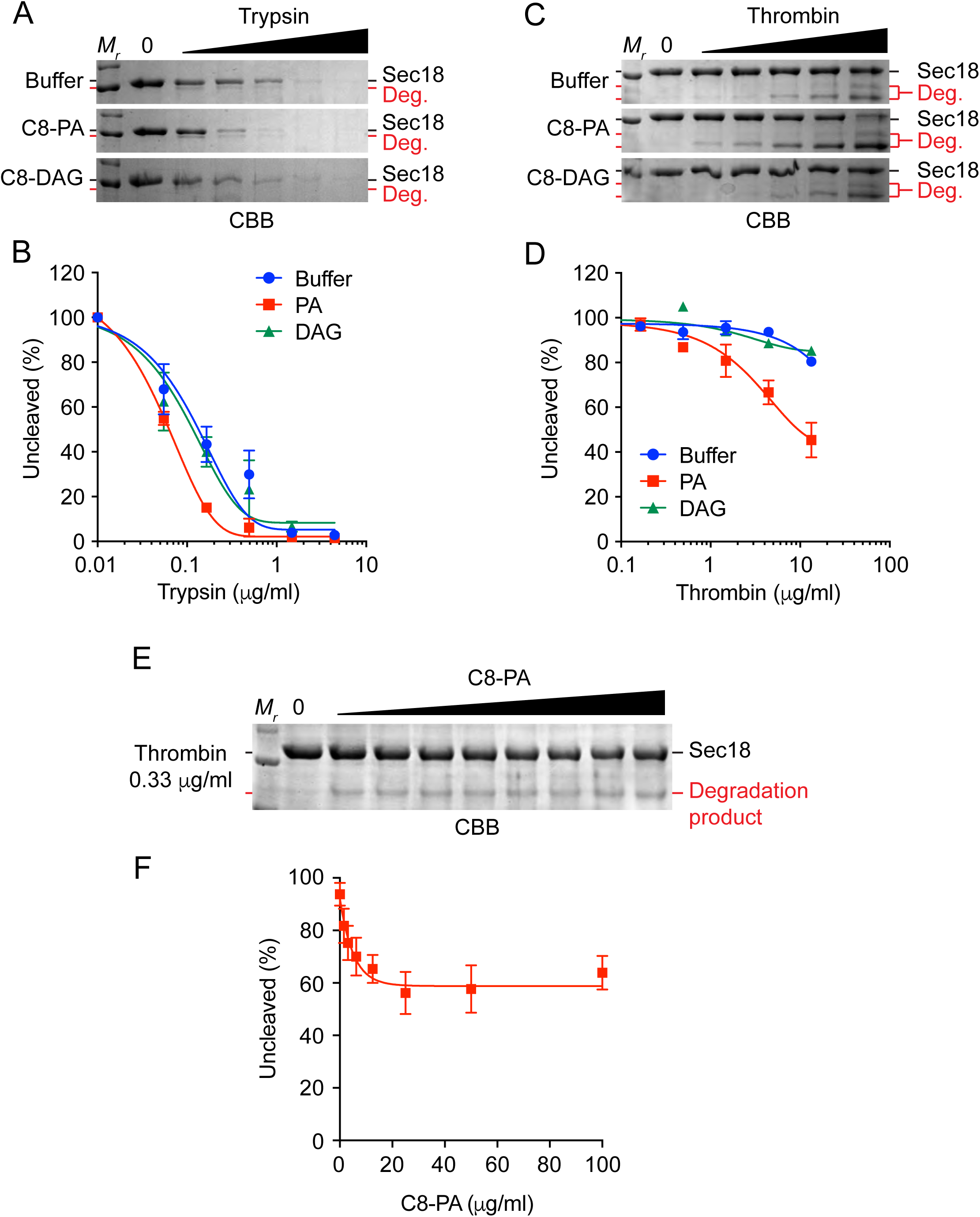
Short chain phosphatidic acid alters the proteolytic cleavage profile of Sec18. Sec18_His8_ was incubated with C8-PA (red), C8-DAG (green), or alone (black) before incubation with increasing concentrations of trypsin (**A**) or thrombin (**C**). Densitometry values of the uncleaved band were measured for each concentration and normalized against the input lane for trypsin (**B**) and thrombin (**D**). Sec18_His8_ was incubated with increasing concentrations of C8-PA before cleavage by thrombin (**E**), and normalized densitometry values against the input control are included (**F**).

Additionally, we performed the same limited proteolysis assay using thrombin in place of trypsin. Thrombin displays much higher specificity than trypsin and should only cleave proteins at specific recognition sites. Incubation of Sec18 with thrombin alone showed no proteolytic degradation of the protein indicating that no recognition sites were accessible to the protease. However, upon addition of C8-PA thrombin was able to cleave Sec18 **(Fig. 4C-D)**. Once again, inclusion of C8-DAG did not show a similar effect indicating once again that the observed conformation change was PA specific. Finally, we titrated C8-PA into a thrombin cleavage assay keeping the concentration of the protease constant.

Cleavage of Sec18 by thrombin showed dose dependence for C8-PA **(Fig. 4E-F)**. These data illustrate that C8-PA binding to Sec18 alters the conformation of the protein allowing for the exposure of an otherwise shielded thrombin recognition site. Sec18 has one predicted thrombin recognition site (after R638) which is located in the D2 domain of the protein [exPASy]. The D2 domain is responsible for the multimerization of Sec18 to its active hexamer when it is in a nucleotide bound state (Lenzen et al., 1998; Yu et al., 1998). This further suggests that PA alters the conformation of the Sec18 D2 domain, or potentially the conformation of D2 with respect to D1 allowing binding to PA. Changes to the D2 domain structure could alter nucleotide binding or disrupt key interactions between protomers thereby decreasing Sec18 hexamer formation. Sec18 is known to associate with cis-SNAREs in its active hexameric form, so inhibition of hexamer formation could decrease its ability to properly recruit to inactive SNARE complexes. This idea is consistent with previous observations that showed increased PA at the vacuole led to decreased recruitment of Sec18 to cis-SNARE complexes (Starr et al., 2016).

### Phosphatidic acid has no significant effect on the secondary structure of Sec18

Because we observed significant changes in the conformation of Sec18 upon binding to C8-PA we next wanted to monitor changes in the secondary structure of the protein when bound to the lipid. To do this we observed the α-helix and β-sheet content of Sec18 in the presence of PA using circular dichroism (CD). CD spectra of mSec18 were obtained in the absence and presence of C8-PA to determine if the protein’s secondary structure was significantly affected by binding the lipid. The spectrum obtained for mSec18 alone showed that the protein was well folded **(Fig. 5A)**. Upon addition of C8-PA, no significant changes were seen in the spectrum suggesting the lipid binding does not alter secondary structure features within the protein.

**Figure 5.**
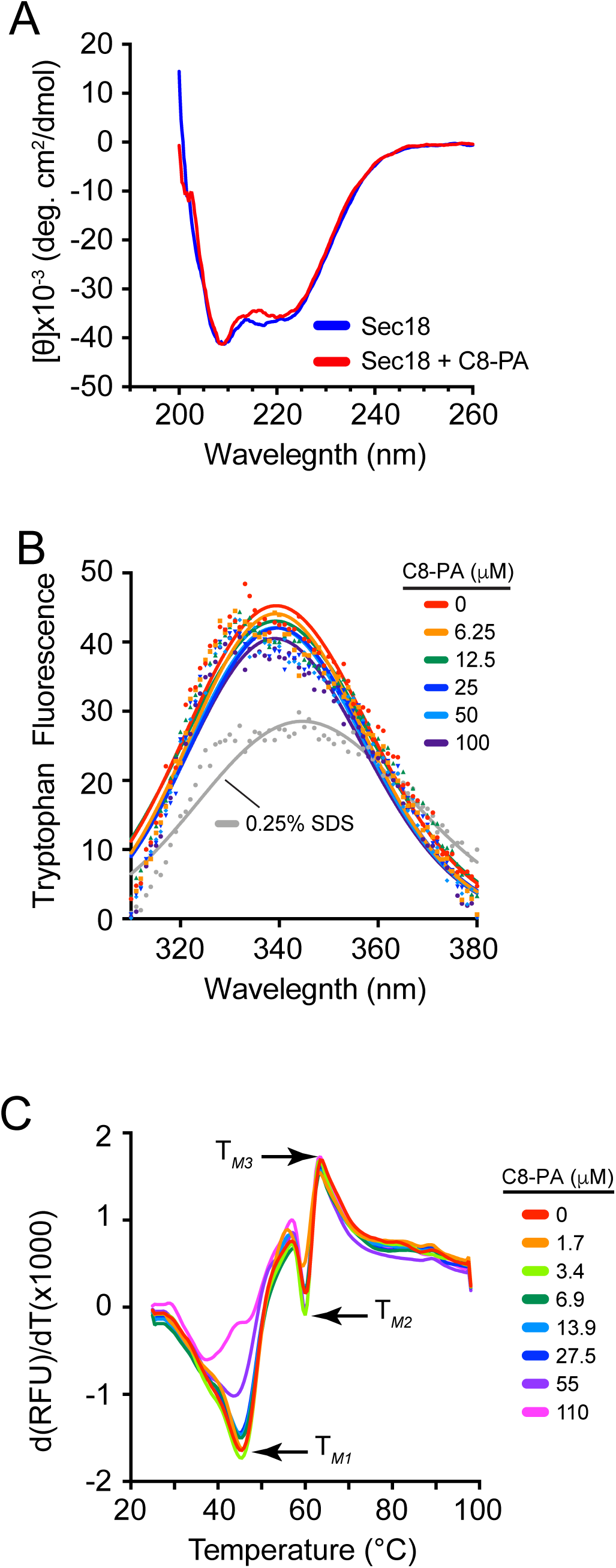
Sec18 does not have significantly altered secondary structure in the presence of short chain phosphatidic acid. (**A**) Circular dichroism spectra were measured (260 to 200 nm, 50 nm min^-1^) for Sec18_His8_ in the presence and absence of C8-PA (100 μM). (**B**) Sec18_His8_ (500 nM) was incubated with increasing concentrations of C8-PA and fluorescence spectra were measured (ex. 295, em. 300-400 nm). The fluorescence (em. 333 nm) for each concentration tested was normalized against the no lipid control and is shown. (**C**) Differential scanning fluorimetry first derivative melting curves were measured (SYPRO orange: ex. 490, em. 560 nm) for increasing concentrations of C8-PA

To rule out any denaturation caused by binding of C8-PA to Sec18, intrinsic tryptophan fluorescence was measured with and without lipid addition. Sec18 contains three tryptophan residues (W88, W91, and W632) in its N and D2 domains. Upon denaturation of Sec18 with SDS, Trp fluorescence was red-shifted and showed decreased intensity **(Fig. 5B)**. Upon incubation with C8-PA, no shift or intensity change was observed. This suggests that PA binding to Sec18 did not lead to denaturation, *i.e*. causing a conformational change large enough to alter the local environment of any of the Trp residues found in the protein.

Finally, to test whether binding PA altered the thermal stability of Sec18, we used differential scanning fluorimetry (DSF) (Miner et al., 2016). Sec18 was labeled with SYPRO orange dye, incubated with different concentrations of C8-PA in separate wells, and equilibrated prior to starting a melting curve. Fluorescence was scanned across a temperature gradient of 20 to 95°C and the first derivative of the fluorescence data was be used to determine the *T*_*m*_ for each condition. DSF has the ability to show multiple melting transitions (Hew et al., 2015; Vollrath et al., 2014). Our data show that mSec18 has three melting transitions. The first mSec18 transition (*T*_M1_) occurred at ~45°C, while *T*_M2_ and *T*_M3_ were at 60°C and 64°C, respectively **(Fig. 5C)**. The addition of C8-PA had no effect on *T*_M2_ and *T*_M3_, as the curves overlapped with the that of apo-Sec18. That said, C8-PA has a striking effect at *T*_*M1*_ where we observed a dose-dependent increase in fluorescence. This likely mirrors the conformational changes seen with limited proteolysis and ANS fluorescence. Taken together these observations lead us to conclude that PA binding to Sec18 induces a significant change to the architecture of the protein but does not denature the protein nonspecifically.

### NSF D1-D2 undergoes large conformational change during transition between hexameric and monomeric forms

To examine the Sec18 conformational changes we observed previously at a more detailed level, atomic molecular dynamics (MD) simulations were performed using NSF, the mammalian homolog of Sec18. The NSF D1-D2 monomer extracted from the cryo-EM structure of an ATP-bound NSF complex (pdb 3J94) after removing bound ATPs was equilibrated with restraints for 20 ns and then relaxed for 200 ns. Based on the overall alpha carbon (*Cα*) RMSD, the monomer undergoes conformational changes up to 15 Å apart from the form originally adopted in the hexamer **(Fig. 6A)**. Calculation on the secondary structure components showed that only the modeled loop region from residue 458 to 478 transitioned from helix during the relaxation to turn and coil (data not shown). This is expected as the loop was poorly resolved in cryo-EM and was only stabilized by interactions with the N-domain in the template crystal structure (the N-D1 domain of p97) used in homology modeling. The stable secondary structure observed in D1 and D2 domains indicated that the large deviation did not come from secondary structural changes, further verifying CD experiments. Instead, we observed that the conformational change was accompanied by an opening-up process of D1 and D2 domains during the relaxation **(Fig. 6A)**. The observation was in agreement with the hypothesis that NSF hexamerization might require certain conformations of D1-D2 monomer and that the conformation required could be further stabilized at the hexamer interface.

**Figure 6.**
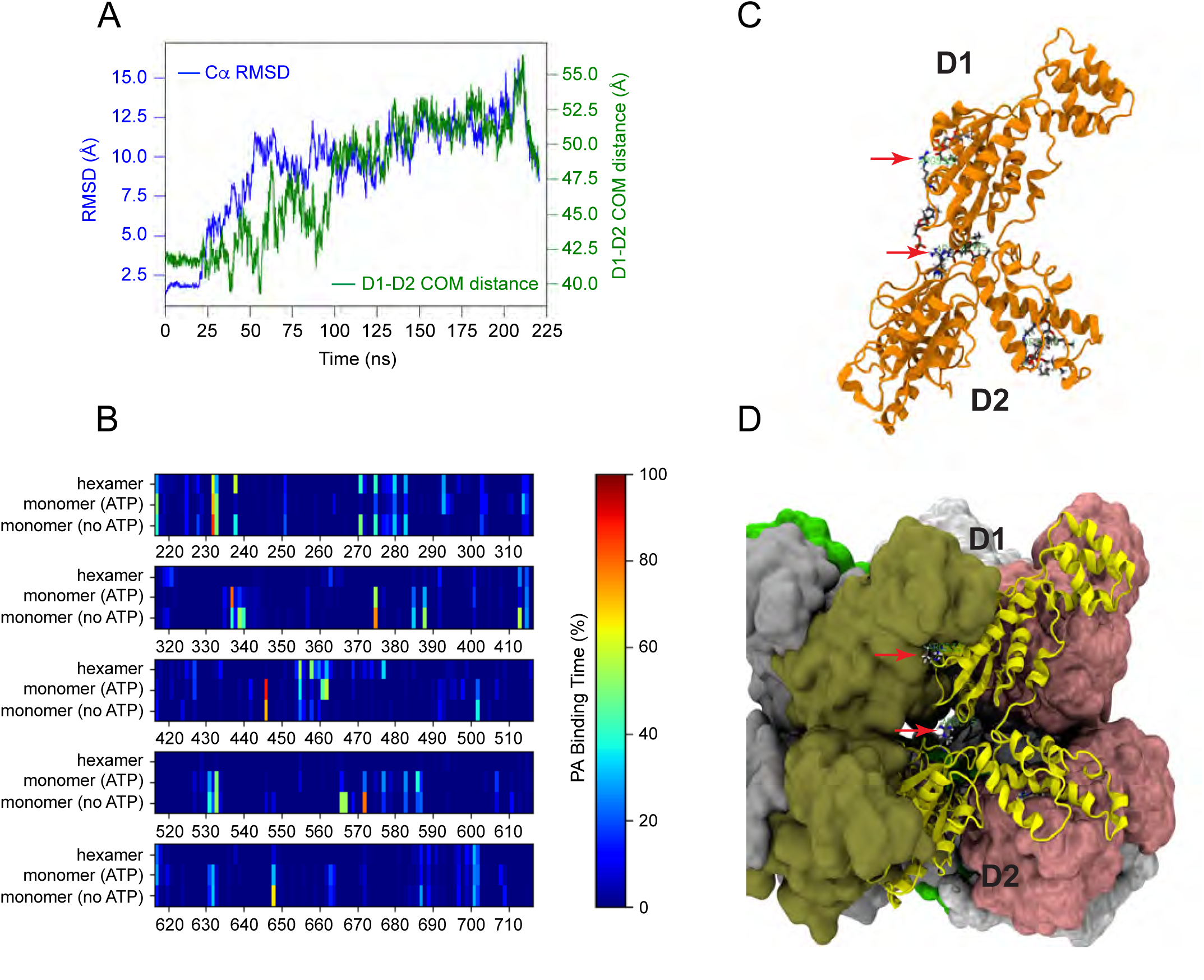
Computational Simulations Show Large Scale Conformational Change Between D1 and D2 Subunits of NSF and Indicate Potential PA binding Regions of NSF. (**A**) D1-D2 monomer undergoes large conformational change during relaxation. In the first 20 ns, D1-D2 monomer was equilibrated with a 0.05 kcal/mol/Å^∧^2 harmonic restraint on protein Ca atoms. Blue: D1-D2 monomer Ca RMSD; green: center of mass distance between D1 and D2 domains. (**B**) Protomer chain A from hexamer cryo-EM structure (PDB: 3J94) was simulated in short-tailed PA solution (119mM, 61 PA molecules in a 95Å x 94Å x120Å water box) for 350 ns with ATP binding and 200 ns without ATP. Binding percentages were measured according to amount of time a PA molecule was within a hydrogen bonding distance from a given amino acid residue of NSF according to heatmap on right side of Figure 6C with residues of NSF indicated on the X axis and model flooded on Y axis. Both monomer (**C**) and hexamer (**D**) are shown with key residues from Fig. 6B indicated on Fig. 6C monomer and Fig. 6D hexamer demonstrating region of hexamer where residues of monomer showing high binding are located.

### Residues of NSF shown to bind to C8PA are not available for PA binding when it is in the hexameric form

Computational flooding studies were performed for both monomeric and hexameric form of the NSF D1-D2 domains, based on the structural information of mammalian NSF **(Fig. 6B)**. Analysis of binding was performed using percent bound as determined by proximity (H-bond distance between phosphate oxygens of PA and between a basic amino acid residue) of PA against time PA ligands were in set proximity. Residues determined to have highest percent bound were determined for both monomeric and hexameric forms of D1-D2. Our flooding simulations of NSF hexamer showed that residues having the highest percent bound PA in the monomer **(Fig. 6C)** were shielded to block lipid binding in the hexameric NSF D1-D2 construct **(Fig. 6D)**. This suggests that PA binding specificity lies somewhere within the hexameric interface.

### Binding Prediction and Clustering Analysis of PA Binding Regions of NSF

Ensemble molecular docking of C8-PA to NSF monomer was performed using the aforementioned D1-D2 equilibrium simulation (Trott *et al*., 2010). Snapshots from the equilibrium trajectory were utilized for molecular docking every 100 ps to fully sample conformational dynamics. The resulting docked C8-PA poses were clustered and an average affinity (Δ*G*) was determined for binding clusters (Beauchamp *et al*., 2011) of monomeric NSF showing similar affinity for C8-PA (~-5 kcal) approximating the MST binding measurements of mSec18 to C8-PA, which is within a ± 2 kcal error prediction ratio used for many docking software programs such as Schrodinger Glide (Friesner et al., 2006) **(Fig. 7A)**. To verify cluster analysis, SiteMap was used and the top 5 site scores taken **(Fig. 7B)**. Figure 7B depicts SiteMap site 1 corresponding to the largest cluster obtained from the ensemble docking **(Fig. 7A)**, which lies in the hexamerization interface illustrated in Figures 7C-D demonstrating the potential importance of this region for PA binding specificity.

**Figure 7.**
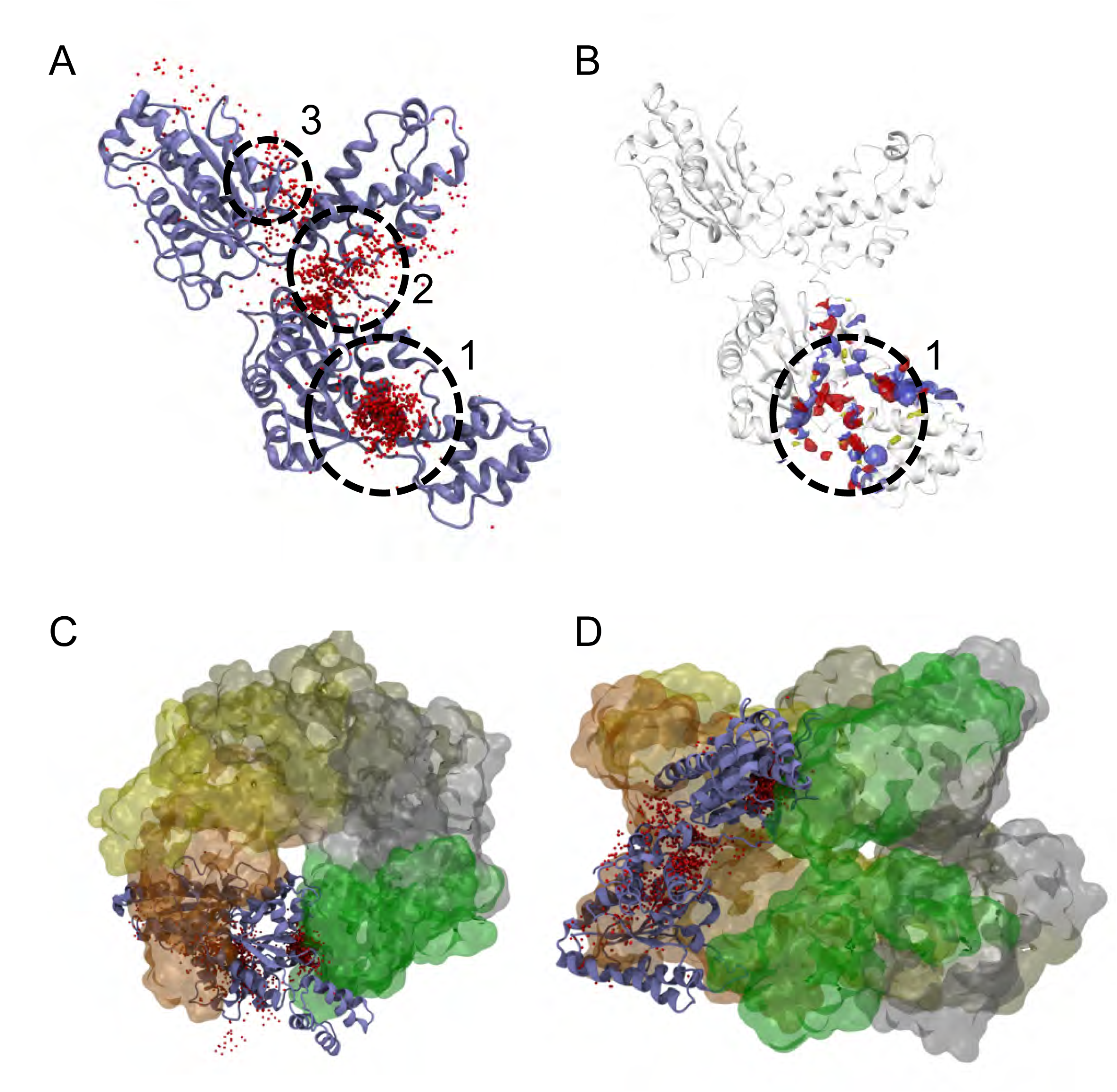
Ensemble Molecular Docking and Binding Site Prediction of NSF D1-D2 Monomer. (**A**) Small red spheres indicate positions on NSF monomer where short chained PA was docked. The circles 1, 2, 3 indicate clusters identified from the top short-chain PA ensemble docking results. (**B**) SiteMap predicted a high affinity binding region of NSF indicated by circle 1, which corresponds to the top ensemble docking cluster from Fig. 7A where yellow indicates potential hydrophobic binding regions, blue indicates potential acidic binding regions, and red indicates potential basic binding regions on NSF. (**C**) A top down D1-D2 depiction of NSF D1-D2 hexamer is shown indicating clusters as described in Fig. 7A are shown relative to the hexamer. (**D**) A side-view D1-D2 depiction of NSF D1-D2 hexamer is shown indicating clusters as described in Fig. 7A are shown relative to the hexamer.

## DISCUSSION

Membrane fusion is a necessary process for all eukaryotes, and Sec18/NSF is the only known protein responsible for utilizing energy from ATP to prime SNAREs (Mayer et al., 1996; Zhao et al., 2015; Ryu et al., 2015). To achieve compartmental specificity, unique SNARE combinations are utilized by defined organelles as well as smaller transport vesicles budding from such organelles (Jahn and Scheller, 2006). Each organelle varies in both size and function, and must contain its own unique combination of protein and lipid factors to allow for specificity in trafficking and membrane fusion events. Regulation of Sec18/NSF is of special significance due its direct role in maintenance of fusion and compartmentalization throughout the eukaryotic cell. Therefore, it is important to understand the role that regulatory factors have on ubiquitous fusion machinery such as Sec18/NSF to adequately model how specificity and efficiency are balanced and maintained at different locations in the cell.

Protein function can be regulated directly through posttranslational modifications or through their interactions with other molecules, including lipids. The vacuole fusion pathway is regulated at various stages by distinct lipids such as phosphoinositides, ergosterol, DAG and PA (Boeddinghaus et al., 2002; Fratti et al., 2004; Jun et al., 2004; Kato and Wickner, 2001; Karunakaran et al., 2012; Karunakaran and Fratti, 2013; Mayer et al., 2000; Miner et al., 2016; Miner et al., 2017; Sasser et al., 2012; Starr et al., 2016; Stroupe et al., 2006). The priming stage requires the presence of ergosterol, PI(4,5)P_2_, as well as the conversion of PA to DAG (Kato and Wickner, 2001; Mayer et al., 2000; Sasser et al., 2012; Starr et al., 2016).

Previously we found that vacuolar PA sequestered Sec18 from *cis*-SNAREs and that the PA phosphatase Pah1/Lipin1 was required to convert PA to DAG to allow Sec18 dissociation from the membrane and recruitment to SNARE complexes (Starr et al., 2016). Although PA turnover is needed for priming, the presence of the lipid is also required downstream for mechanisms that remain to be characterized. Deletion of *PAH1* or the DAG kinase *DGK1* alters the balance of PA and DAG on vacuole to dramatically affect membrane fusion (Sasser et al., 2012; Miner et al., 2017). We thus postulate that enzymatic changes that alter PA levels can in turn shift the equilibrium of Sec18 from a lipid-bound to a to a SNARE-associated state. Such changes would likely have significant effects on SNARE activation and the overall progression of the membrane fusion cascade.

In this study we demonstrated that Sec18 directly binds PA with high affinity on par with a known PA-binding domain. Moreover, only monomeric Sec18 could bind both PA membranes and soluble C8-PA, whereas hexameric was only able to bind C8-PA. This signifies that C8-PA could access PA-binding residues that are blocked in the hexamer to prevent membrane association. Sec18/NSF exists as both a monomer (non-enzymatic) and a hexamer (enzymatic). Our findings indicate that Sec18 exists in both a monomeric lipid-bound pool and SNARE-bound hexamers. Because ATP is required for Sec18 hexamerization, we tested PA liposome binding at two concentrations of ATP. The D1 domain of NSF has *K*_*D*_ of ~20 μM for ATP, while the D2 hexamerization domain has a *K*_*D*_ of ~40 nM. PA binding was unaffected 100 nM ATP that should predominantly bind D2; however, at 100 μM ATP mSec18 binding to PA was abolished. This suggests that the nucleotide bound state of the D1 domain is important in specific PA binding of Sec18/NSF, likely due to conformational changes in the ATP bound state. Alternatively, the higher ATP concentration may shift the monomeric pool of Sec18 used to a predominantly hexameric pool decreasing its affinity for PA.

During priming, Sec17/α-Snap is recognized by Sec18/NSF in an ATP bound state at D1 before subsequent ATPase activity occurs. We posit that Sec18 exists in both lipid-bound and SNARE-bound states and that the presence of ATP at the D1 NBD may determine the state in which the protein primarily exists. Membrane PA may prevent the association of ATP with the D1 NBD locking the protein in an inactive lipid-bound state preventing recruitment to inactive SNARE complexes. This is in line with our data in this study and with observations from previous work (Starr et al., 2016).

The fact that Sec18 monomer binding to PA liposomes was inhibited at a saturating ATP concentration for the D1 NBD could indicate that the PA binding site for Sec18 lies near the D1 ATP binding site. Alternatively, it is possible the conformation of Sec18 in its ATP bound state shields the protein’s unique PA binding site. The idea that Sec18 binding to PA may not specifically depend on the D1 ATP binding site was supported by computational flooding experiments performed on both hexamer and monomer in the presence and absence of ATP. Flooding experiments allowed for C8-PA to equilibrate with NSF monomer, and binding was measured using the length of time a PA molecule resided near a given residue of NSF. Many of the long term amino acid residues sharing the longest contact time to PA were predictably basic residues, especially lysine and arginine. However, dramatic differences in these residues were not noticed between the ATP and non-ATP simulations. Furthermore, many of the residues with longer PA binding time were not of importance for PA binding in the hexamer simulation. This result is in corroboration with the high binding affinity of Sec18 monomers to PA liposomes vs the hexameric form. This further indicates that the Sec18 monomer and hexamer are differentially regulated. Furthermore, it suggests that PA may influence the formation of the active hexamer by controlling the availability of its inactive monomer at membranes.

We propose that Sec18/NSF PA regulation is achieved by sequestration of Sec18/NSF monomer on PA containing membranes blocking its ability to form active hexamer. This sequestration regulates the ability of Sec18 to perform its enzymatic function of activating SNAREs. Additionally, it is possible that PA at the site of priming could increase localization of Sec18/NSF to the membrane in preparation of SNARE priming. Additional factors, such as the PA phosphatase Pah1/Lipin, could serve to activate Sec18/NSF once the fusion cascade was required to proceed (Sasser et al., 2012; Starr et al., 2016). In this way, PA could serve as a temporal regulator of SNARE priming activity and of the membrane fusion process as a whole.

Two additional modes of regulation are suggested by our data. First, PA membrane concentrations may play either a local or organelle-dependent role for the regulation of Sec18/NSF by lipid binding. Depending on the concentration and localization of PA at a given membrane Sec18 sequestration by PA could either play a larger or less prominent role in regulating the priming of SNAREs. Second, we have found that curvature plays a role in Sec18 regulation by PA. Therefore, Sec18 regulation by PA could be differentially managed in the cell based on vesicle or organelle size.

This work has shown that Sec18/NSF binds to membranes containing PA with high affinity and that this lipid binding is greater to the monomeric version of the protein. Upon binding PA, Sec18/NSF undergoes a significant conformational change that coincides with a reduction in the protein’s SNARE priming activity. Molecular dynamics simulations show that monomeric Sec18/NSF has greater conformational flexibility than hexamer. Equilibrium simulations indicate a large scale conformational change when NSF hexamer is converted to monomer **(Fig. 6B)**. We propose that binding to PA by Sec18/NSF may serve a regulatory role in preventing the formation of the active hexamer thereby throttling the priming of SNAREs. There are two main reasons we think this form of regulation is needed: 1) To keep NSF from freely priming any cis-SNARE complex around (Yavuz et al., 2018), and 2) To keep NSF nearby without allowing priming.

We have identified regions of NSF that are potential high PA binders **(Fig. 7A-B)**. Additionally, we have identified residues of NSF that show potential PA binding in monomeric form that are not capable of binding PA in hexameric form **(Fig. 6B)**. In order for Sec18 to form hexamer, according to our model, Sec18 would have to be removed from the membrane, and we have previously shown that the only PA phosphatase that plays a role in membrane fusion is Pah1 (Sasser et al., 2012). It is possible that an additional chaperone may be involved in alleviating the transition of Sec18 monomer towards hexamerization.

Based on our computational studies, it appears that there are numerous candidate residues that might contribute to Sec18 PA binding. HMMM simulations have been performed (data not shown); however, due to the size and flexibility of Sec18 monomer, long time scales in the micro second range may be required to show final binding events sequestering Sec18 to a PA containing membrane. We plan to further probe this binding event using HMMM at a longer time scale to capture the exact binding event of Sec18 to a PA membrane, and to specifically identify the numerous residues that may be involved.

## MATERIALS AND METHODS

### Reagents

POPA (1-palmitoyl-2-oleoyl-sn-glycero-3-phosphate), POPC (1-palmitoyl-2-oleoyl-sn-glycero-3-phosphatidylcholine), POPE (1-palmitoyl-2-oleoyl-sn-glycero-3-phosphatidylethanolamine), C8-PA (1,2-dioctanoyl-sn-glycero-3-phosphate), C8-DAG (1,2-dioctanoyl-sn-glycerol), and C8-PS (1,2-dioctanoyl-sn-glycero-3-phospho-L-serine) were purchased from Avanti Polar Lipids (Alabaster, AL, USA) as chloroform stock solutions and stored at −20°C. CM7 and Ni-NTA (Standard and S series) sensor chips, and Regeneration buffers (Glycine pH 1-3) were procured from GE Healthcare (Buckinghamshire UK). Ni-NTA Atto 488 dye was procured from Sigma-Aldrich Corp. (St. Louis Missouri). Monolith NT.115 standard treated capillaries for thermophoresis were purchased from Nanotemper (München Germany).

### Plasmid construction

Plasmid for expression of Sec18_His8_ was created by amplification of *SEC18* by PCR from genomic DNA of the yeast strain DKY6281 using primers containing Ndel and Xhol restriction cut sites (Forward: 5’-ACGTACGTCAWGTTCAAGATACCTGGTTTTGG-3’, Reverse: 5’-ATCGAATGCTCGAGT-GCGGATTGGGTCAT CAACT-3’). PCR Product was inserted into pET42a using Ndel and Xhol in frame with a C-terminal 8xHis tag sequence under the control of a T7 promoter to create pSec18H8.

Plasmid for expression of GST-Sec18 was created using primers containing EcoRI and XhoI restriction cut sites (Forward: 5’-ATGCAATGGAATTCATGTTCAAGATACCTGGTTTTGG-3’, Reverse: 5’-ATCGAATGCTC GAGTTATGCGGATTGGGTCATCAACT-3’). PCR product was inserted into pParallel GST using EcoRI and XhoI to create pGSTSec18. Plasmid for expression of GST-N terminal domain was created in the same way using a different reverse primer (Forward: 5’-ATGCAATGGAATTCATGTTCAAGATACCTGGTTTTGG-3’, Reverse: 5’-ATCGAATGCTCGAGTCTTCCTTTGAAAAAATTAATTTGTGTTTGTTT-3’) to create pGSTN.

### Protein purification

For purification, pSec18His8 was transformed into Rosetta 2 (DE3) pLysS Competent Cells (Novagen) and Sec18_His8_ expression was carried out using auto-inducing medium (AIM) (Studier, 2014). Cells were grown in AIM until reaching stationary phase (37°C, 18 hours, shaking) and harvested by centrifugation. Cells were resuspended in lysis buffer (20 mM HEPES pH=6.8, 300 mM NaCl, 0.1% Triton-100, 2 mM 2-mercaptoethanol, 20 mM imidazole, 10% glycerol, 1 mM ATP, 1 mM PMSF, and 1X cOmplete Protease Inhibitor Cocktail (Roche)) and lysed by French press. Lysates were cleared by centrifugation (50,000 *× g*, 20 min, 4°C) and incubated with Ni-NTA resin (Invitrogen) overnight at 4°C. Resin was washed with 100 bed volumes of wash buffer (lysis buffer with 50 mM imidazole) before protein was eluted in 1 ml fractions (lysis buffer with 250 mM imidazole). Protein was concentrated before being run through gel filtration (Superose 6) using size exclusion buffer (20 mM HEPES pH 6.8, 300mM NaCl, 1 mM 2-mercaptoethanol, 10% glycerol). Sec18His8 elutes in two peaks corresponding to monomeric and hexameric pools. Each pool was collected and concentrated before use. For circular dichroism experiments, Sec18_His8_ was purified using the same approach with different buffer compositions. CD lysis buffer (50 mM phosphate buffer pH 6.8, 20 mM imidazole, 1mM PMSF), CD wash buffer (50 mM phosphate buffer pH 6.8, 50 mM imidazole), CD elution buffer (50 mM phosphate buffer pH 6.8, 250 mM imidazole), and CD SEC buffer (50 mM phosphate buffer pH 6.8) were used. GST-Sec18 was purified similarly using Rosetta 2 (DE3) pLysS Competent Cells transformed with pGSTSec18 but with the following changes. GST lysis buffer (50 mM Tris pH 8.0, 150 mM NaCl, 5 mM EDTA, 1 mM ATP, 1 mM PMSF, and 1X cOmplete Protease Inhibitor Cocktail) was used through the lysis and chromatography wash steps. Protein was eluted with GST elution buffer (20 mM HEPES pH 7.2, 150 mM NaCl, 10 mM reduced glutathione) and dialyzed against 1X HBS pH 7.2 before being aliquoted and stored at −80°C. GST-N was purified in the same way using cells transformed with pGST-N. The DEP PA binding domain from murine Dvl2 was purified as a GST-fusion as described (Capelluto et al., 2014). Membrane scaffold protein 1D1 (MSP1D1-His) was prepared as described (Denisov et al., 2004).

### Nanodisc Preparation

Lipid composition of PA nanodiscs consisting of 3.023 μmol POPC diC16, .098 μmol PA diC16, and .78 μmol POPE and PC nanodiscs consisting of 3.121 μmol POPC diC16 and .78 μmol POPE were combined, dried, and desiccated overnight. Lipids were then dissolved in 20 mM sodium deoxycholate in TBS (50 mM Tris-HCl, pH 7.4, 150 mM NaCl, and .02% NaN_3_) and sonicated. MSP1D1 membrane scaffold protein (MSP) was then added in a ratio of 70:1 lipid to protein and detergent removed with Bio-Beads^®^ SM-2 (Bio-Rad). Nanodiscs were isolated using size exclusion chromatograph and quantified using a NanoDrop and the extinction coefficient of 21,000 L mol^-1^ cm^-1^ for MSP1D1 (24.66 kD), and the resultant mg/mL divided by two because there are two MSP proteins per nanodisc (Sparks et al., 2016).

### Surface Plasmon Resonance

Surface plasmon resonance (SPR) measurements were performed on a Biacore T200 instrument equipped with an Ni-NTA chip. Approximately 2000 RU of 5% PA nanodiscs were immobilized non-covalently using 100 mM NiSO4 flowed at 10 μL/s followed by a blank buffer injection of HEPES pH 7.4, 150 mM NaCl (HBS Buffer). Injections were performed in HBS buffer at a flow rate of 30 μl/min with an association time of 90 sec, dissociation time of 300 sec., and binding was measured in relative response units (RU) as described (Sparks et al., 2016). Regeneration with EDTA was performed at flow rate 30 μL/s for 120 s using 100 μM EDTA buffer. Proteins were injected using 1:1 dilutions from highest concentration and steady state was obtained using GE BIAcore T200 evaluation software version 3.0 (BIAevaluate). Proteins were injected using 1:1 dilutions for Sec 18 monomer (3.64 μM, 1,82 μM, 911 nM, and 455 nM), DEP PA binding domain (57.5 μM, 28.8 μM, 14.4 μM, 7.2 μM, 3.6 μM, 5.8 μM), and N domain from Sec18 (84.3 μM, 8.4 μM, 4.2 μM, 1.1 μM, 527 nM, and 1.69 μM) with one concentration from each titration run in duplicate. Steady state data was fitted and exported using BiaEvaluate software into GraphPad Prism 7.00 for Windows, GraphPad Software (La Jolla, CA).

### Microscale thermophoresis

Thermophoresis measurements were performed using a Monolith NT.115 labeled thermophoresis machine. Sec18_His8_ was labeled with Ni-NTA Atto 488 according to the manufacturer’s protocol. M.O. Control software was used for operation of MST. Target protein concentrations were 50 nM for all His-tag labeled proteins Sec 18 monomer, Sec18 hexamer, PA nanodiscs, and PC nanodiscs. LED excitation power was set to 90% and MST set to high allowing 3 seconds prior to MST on to check for initial fluorescence differences, 25 s for thermophoresis, and 3 s for regeneration after MST off. Analysis was performed using M.O. Affinity Analysis Software as the difference between initial fluorescence measure in the first 5 s as compared with thermophoresis at 15 s. All measurements were performed in PBS buffer (137 mM NaCl, 2.7 mM KCl, 8 mM Na2HPO4, and 2 mM KH2PO4, pH 7.4) without Tween except for Sec18 Hexamer, which was performed in 50% PBS buffer and 50% Storage buffer (20 mM HEPES pH=6.8 300 mM NaCl 1 mM beta-mercaptoethanol 10% glycerol) and binding affinity was generated using Graphpad Sigmoidal 4PL fit from points exported from M.O. Affinity Analysis software using *K*_*D*_ Model with target concentration fixed at 50 nM generating bound and unbound, and fraction bound data exported to Graphpad to generate Figures using standard curve Sigmoidal for final *K*_*D*_.

### Limited Proteolysis

Cleavage reactions were carried out in proteolysis buffer (20 mM HEPES pH 7.2, 150 mM NaCl, 2 mM ATP, 2 mM MgCl_2_). Sec18_His8_ (2 μM) was added to proteolysis buffer and incubated with indicated lipid concentration on ice for 5 min. Trypsin or thrombin diluted in 1X HBS was added to assay tubes at indicated concentrations and incubated at 25°C for 30 min. Cleavage reactions were stopped with the addition of SDS sample buffer containing 1 mM PMSF. Samples were resolved with SDS-PAGE and gels were stained using Coomassie Blue. Gels were destained with methanol/acetic acid solution (50%/7%) and imaged using a ChemiDoc MP Imaging System (Bio-Rad).

### Tryptophan Fluorescence Spectroscopy

Sec18_His8_ (500 nM) was incubated with the indicated concentrations of C8-PA in fluorescence assay buffer (20 mM HEPES pH 7.2, 150 mM NaCl, 1 mM MgCl_2_, 1 mM ATP). Lipid dilutions were first prepared in assay buffer and measured for background fluorescence before Sec18_His8_ was added and incubated at 25°C. Intrinsic tryptophan fluorescence measurements were made using a fluorimeter with Peltier temperature control (Agilent Technologies). Samples were excited at 295 nm and the emission spectra were collected from 300-400 nm. Samples were measured in a 100 μL cuvette (Starna Cells). Initial background fluorescence spectra for each lipid concentration were subtracted from final measurements.

### 1,8-ANS Fluorescence Spectroscopy

ANS binding experiments were carried out in fluorescence assay buffer with 5 μM 1-anilino-8-naphthalenesulfonate (ANS) (Cayman Chemical). Initial spectra were taken without Sec18_His8_ to measure any background fluorescence from buffer or added lipids (ex. 350 nm, em. 390-620 nm). Sec18_His8_ diluted in assay conditions was then added to the assay to the indicated concentration and incubated at 25°C for 5 min before spectra were obtained. Initial background fluorescence spectra for each lipid concentration were subtracted from final measurements.

### Circular Dichroism

Monomeric Sec18_His8_ purified in phosphate buffer was incubated with and without C8-PA to equilibrium (25°C, 15 min). Protein concentration used was 5 μM and lipid concentration used was 100 μM. Circular dichroism was measured using a spectropolarimeter (JASCO). All spectra were recorded from 260 nm to 200 nm at 50 nm min^-1^ and measurements were taken in a 1 mm pathlength cuvette.

### Differential Scanning Fluorimetry

Sec18 (2.75 mg/mL) was diluted to a final concentration of 0.11 mg/mL in phosphate buffer containing 1 mM ATP, 1 mM MgCl_2_, and 4X SYPRO orange dye. Next, 22.5 μL of this mix was added to a white hardshell 96-well PCR plate (Bio-Rad) which contained 2.5 μL of serial dilutions of C8-PA in phosphate buffer. The plates were then sealed with Microseal ‘B’ film (Bio-Rad), and samples were allowed to equilibrate at room temperature for 30 min before beginning the assay. Melting curves were performed using a Bio-Rad CFX Connect real-time detection system. The melt curve protocol was 25°C for 3 min followed by a 25-90°C gradient with 0.5°C increments. Each temperature was held for 10 seconds and the fluorescence intensity was measured (Ex = 490 nm, Em = 560 nm). The first derivative of the fluorescence readings was used to determine the melting temperature(s) for each condition.

### Preparation of D1-D2 monomer and hexamer models

The D1-D2 monomer model (residues 215-737) was derived from an Cryo-EM structure of ATP-bound NSF complex (PDB 3J94 - chain A) (Zhao et al., 2015). Missing residues [335-346, 458-478 in PDB 3J94 (chain A)] were built via homology modeling using the crystal structure of the homologous N-D1 domain of p97 (PDB 1E32) as a template by MODELLER 9.19 (Sali and Blundell, 1993). The complete D1-D2 hexamer model was prepared (Jo et al., 2014) using the same PDB 3J94 as the monomer. Missing loops in each monomer were modeled in CHARMM GUI to ensure that no clashes or topological errors exist in the complex structure. Cis-peptide bonds in both monomer and hexamer structures were examined and fixed manually using Cispeptide plugin in VMD (Schreiner et al., 2011). A further refinement of loops built in the hexamer was performed via MDFF (Trabuco et al., 2008).

### Equilibrium MD simulations of D1-D2 monomer and D1-D2 hexamer

The MD simulations were performed with NAMD 2.12 (Phillips et al., 2005) using CHARMM36m force field (Huang et al., 2017). Langevin dynamics and Langevin piston Nosé-Hoover methods (Feller et al., 1995; Martyna et al., 1994) were used to maintain constant temperature at 310.15 K and pressure at 1 atm. The long-range electrostatic forces were evaluated using the particle mesh Ewald (PME) method (Darden et al., 1993; Essmann et al., 1995) with a 1 Å grid spacing. The van der Waals interactions were calculated with a cutoff of 12 Å and a force-based switching scheme after 10 Å. Integration time step was set at 2 fs with SETTLE algorithm (Miyamoto and Kollman, 1992) applied. VMD 1.9.3 was used for MD trajectory visualization and analysis (Humphrey et al., 1996). The D1-D2 monomer model was first equilibrated for 20 ns with harmonic restraints (0.05 kcal/mol/Å^2^) on protein Cα atoms except modelled loops, then followed by 200 ns equilibration without restraints.

### PA lipids flooding simulations of D1-D2 monomer and D1-D2 hexamer

All the structures of D1-D2 monomer and hexamer were combined with a lipid grid of 5 × 5 × 5 short-chain PA lipids consisting of protein, solvent, and lipid with overlapping lipids on protein removed. The flooding box was then solvated and ionized with the SOLVATE and AUTOIONIZE plugins within VMD [PMID: 8744570] with a final NaCl concentration of 150 mM. The constructed D1-D2 monomer (ATP-bound and ATP-free) and D1-D2 hexamer (ATP-bound) were simulated in a short-chain PA solution for 120 ns each for monomer and 166 ns for hexamer, termed 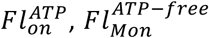 and 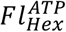. Final simulation systems include for monomer ~80 mM PA lipids in water and for hexamer short-chain PA solution (120 mM, 223 PA molecules in a 188 Å × 187 Å × 133 Å water box) for 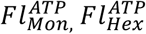, and 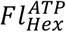. Harmonic restraints were applied on protein C_α_ atoms except modelled loops throughout the simulation, to preserve protein secondary structure.

### Binding Site Simulations on NSF for PA

To characterize C8-PA and D1-D2 monomer interactions, molecular ensemble docking of PA was done on D1-D2 monomer using AutoDock Vina (Pande et al., 2010). The previously mentioned equilibrium simulation of D1-D2 was used to fully sample the dynamics of D1-D2 for molecular docking, where snapshots were taken every 100 ps of the 200 ns trajectory. For each snapshot, an 80Å by 94Å by 108Å grid box was used to fully sample the entire structure. Each snapshot was docked with an exhaustiveness of 10, yielding a total of 2000 PA docked poses, with the affinities of each poses obtained from the resultant log files. These poses where then clustered using a hybrid K-centers and K-medoids clustering algorithm using root-mean-square deviation (RMSD) method, (Beauchamp et al., 2011) for which three main clusters where identified. These clusters where then compared to SiteMap (Halgren, 2009). Schrodinger SiteMap was used on equilibrated D1-D2 NSF monomer indicating top potential ligand binding regions of NSF D1-D2 monomer including shallow binding sites.

### Statistics

Results are expressed as the mean ± S.E. where n = number of replicates. SPR experiments were analyzed using GE BIAevaluation Software, MST experiments were analyzed using M.O. Affinity Analysis software and GraphPad Prism was utilized for statistical approaches.

## Acknowledgements

This research was supported by grants from the National Institutes of Health (R01-GM101132 to RAF, and P41-GM104601, U01-GM111251 and U54-GM087519 to ET), the National Science Foundation (MCB 1818310 to RAF), and the Office of Naval Research (ONR N00014-16-1-2535 to E.T.). Computational resources were provided by XSEDE (XSEDE MCA06N060) and Blue Waters (ACI-1440026). SPR was aided by the help of Dr. Jermaine Jenkins at the University of Rochester Structural Biology & Biophysics Facility with support from NIH NCRR grant 1S10 RR027241, as well as NIH NIAID P30AI078498 and the University of Rochester School of Medicine and Dentistry. We also thank Dr. Susan Martinis and Mr. Zach Dayu for help preparing nanodiscs.

## Author Contributions

MLS, RPS, LRH, JLJ, ZZ, AA, ML and RAF designed research; MLS, RPS, LHR, JLJ, ZZ, AA, and ML performed research; MLS, RPS, LRH, JLJ, ZZ, AA, ML, ET and RAF analyzed data; MLS, RPS, and RAF wrote the paper.

## Abbreviations

diC8: dioctanoyl
PA: phosphatidic acid
DAG: diacylglycerol
PC: phosphatidylcholine
PE: phosphatidylethanolamine
PS: phosphatidylserine
MSP: membrane scaffold protein
NEM: N-ethylmaleimide
NSF: NEM sensitive factor
α-SNAP: soluble NSF adaptor protein
SNARE: soluble N-ethylmaleimide-sensitive factor attachment protein receptor

## REFERENCES

1. Beauchamp, K. A., Bowman, G. R., Lane, T. J., Maibaum, L., Haque, I. S., and Pande, V. S. (2011). MSMBuilder2: Modeling Conformational Dynamics at the Picosecond to Millisecond Scale. J Chem Theory Comput 7, 3412–3419.

2. Boeddinghaus, C., Merz, A. J., Laage, R., and Ungermann, C. (2002). A cycle of Vam7p release from and PtdIns 3-P-dependent rebinding to the yeast vacuole is required for homotypic vacuole fusion. J Cell Biol 157, 79–89.

3. Capelluto, D. G., Zhao, X., Lucas, A., Lemkul, J. A., Xiao, S., Fu, X., Sun, F., Bevan, D. R., and Finkielstein, C. V. (2014). Biophysical and molecular-dynamics studies of phosphatidic acid binding by the Dvl-2 DEP domain. Biophys J 106, 1101–1111.

4. Chang, L. F., Chen, S., Liu, C. C., Pan, X., Jiang, J., Bai, X. C., Xie, X., Wang, H. W., and Sui, S. F. (2012). Structural characterization of full-length NSF and 20S particles. Nat Struct Mol Biol 19, 268–275.

5. Darden, T., York, D., and Pedersen, L. G. (1993). Particle Mesh Ewald: An N·log(N) Method for Ewald Sums in Large Systems. J Chem Phys 98, 10089–10092.

6. Denisov, I. G., Grinkova, Y. V., Lazarides, A. A., and Sligar, S. G. (2004). Directed self-assembly of monodisperse phospholipid bilayer Nanodiscs with controlled size. J Am Chem Soc 126, 3477–3487.

7. Essmann, U., Perera, L., Berkowitz, M. L., Darden, T., Lee, H., and Pedersen, L. G. (1995). A Smooth Particle Mesh Ewald: An N-log(N) Method for Ewald Sums in Large Systems. J Chem Phys 103, 8577–8593.

8. Feller, S. E., Zhang, Y., Pastor, R. W., and Brooks, B. R. (1995). Constant Pressure Molecular Dynamics Simulation: The Langevin Piston Method. J Chem Phys 103, 4613–4621.

9. Fleming, K. G., Hohl, T. M., Yu, R. C., Müller, S. A., Wolpensinger, B., Engel, A., Engelhardt, H., Brünger, A. T., Söllner, T. H., and Hanson, P. I. (1998). A revised model for the oligomeric state of the N-ethylmaleimide-sensitive fusion protein, NSF. J Biol Chem 273, 15675–15681.

10. Fratti, R. A., Jun, Y., Merz, A. J., Margolis, N., and Wickner, W. (2004). Interdependent assembly of specific regulatory lipids and membrane fusion proteins into the vertex ring domain of docked vacuoles. J Cell Biol 167, 1087–1098.

11. Friesner, R. A., Murphy, R. B., Repasky, M. P., Frye, L. L., Greenwood, J. R., Halgren, T. A., Sanschagrin, P. C., and Mainz, D. T. (2006). Extra precision glide: docking and scoring incorporating a model of hydrophobic enclosure for protein-ligand complexes. J Med Chem 49, 6177–6196.

12. Halgren, T. A. (2009). Identifying and characterizing binding sites and assessing druggability. J Chem Inf Model 49, 377–389.

13. Hew, K., Dahlroth, S. L., Veerappan, S., Pan, L. X., Cornvik, T., and Nordlund, P. (2015). Structure of the Varicella Zoster Virus Thymidylate Synthase Establishes Functional and Structural Similarities as the Human Enzyme and Potentiates Itself as a Target of Brivudine. PLoS One 10, e0143947.

14. Heyduk, T., and Lee, J. C. (1989). Escherichia coli cAMP receptor protein: evidence for three protein conformational states with different promoter binding affinities. Biochemistry 28, 6914–6924.

15. Huang, J., Rauscher, S., Nawrocki, G., Ran, T., Feig, M., de Groot, B. L., Grubmüller, H., and MacKerell, A. D. (2017). CHARMM36m: an improved force field for folded and intrinsically disordered proteins. Nat Methods 14, 71–73.

16. Humphrey, W., Dalke, A., and Schulten, K. (1996). VMD - Visual Molecular Dynamics. J Mol Graphics 14, 22–28.

17. Jahn, R., Lang, T., and Südhof, T. C. (2003). Membrane fusion. Cell 112, 519–533.

18. Jahn, R., and Scheller, R. H. (2006). SNAREs - engines for membrane fusion. Nat Rev Mol Cell Biol 7, 631–643.

19. Jahn, R., and Sudhof, T. C. (1999). Membrane fusion and exocytosis. Annu Rev Biochem 68, 863–911.

20. Jo, S., Cheng, X., Islam, S. M., Huang, L., Rui, H., Zhu, A., Lee, H. S., Qi, Y., Han, W., Vanommeslaeghe, K., MacKerell, A. D., Roux, B., and Im, W. (2014). CHARMM-GUI PDB manipulator for advanced modeling and simulations of proteins containing nonstandard residues. Adv Protein Chem Struct Biol 96, 235–265.

21. Jun, Y., Fratti, R. A., and Wickner, W. (2004). Diacylglycerol and its formation by Phospholipase C regulate Rab‐ and SNARE-dependent yeast vacuole fusion. J Biol Chem 279, 53186–53195.

22. Karunakaran, S., and Fratti, R. (2013). The Lipid Composition and Physical Properties of the Yeast Vacuole Affect the Hemifusion-Fusion Transition. Traffic 14, 650–662.

23. Karunakaran, S., Sasser, T., Rajalekshmi, S., and Fratti, R. A. (2012). SNAREs, HOPS, and regulatory lipids control the dynamics of vacuolar actin during homotypic fusion. J Cell Sci 14, 1683–1692.

24. Kato, M., and Wickner, W. (2001). Ergosterol is required for the Sec18/ATP-dependent priming step of homotypic vacuole fusion. Embo J 20, 4035–40.

25. Lapinski, M. M., Castro-Forero, A., Greiner, A. J., Ofoli, R. Y., and Blanchard, G. J. (2007). Comparison of liposomes formed by sonication and extrusion: rotational and translational diffusion of an embedded chromophore. Langmuir 23, 11677–11683.

26. Lenzen, C. U., Steinmann, D., Whiteheart, S. W., and Weis, W. I. (1998). Crystal structure of the hexamerization domain of N-ethylmaleimide-sensitive fusion protein. Cell 94, 525–536.

27. Liu, S., Wilson, K. A., Rice-Stitt, T., Neiman, A. M., and McNew, J. A. (2007). In vitro fusion catalyzed by the sporulation-specific t-SNARE light-chain Spo20p is stimulated by phosphatidic acid. Traffic 8, 1630–1643.

28. Manifava, M., Thuring, J. W., Lim, Z. Y., Packman, L., Holmes, A. B., and Ktistakis, N. T. (2001). Differential binding of traffic-related proteins to phosphatidic acid-or phosphatidylinositol (4,5)-bisphosphate-coupled affinity reagents. J Biol Chem 276, 8987–8994.

29. Martyna, G. J., Tobias, D. J., and Klein, M. L. (1994). Constant Pressure Molecular Dynamics Algorithms. J Chem Phys 101, 4177–4189.

30. Matveeva, E. A., He, P., and Whiteheart, S. W. (1997). N-Ethylmaleimide-sensitive fusion protein contains high and low affinity ATP-binding sites that are functionally distinct. J Biol Chem 272, 26413–26418.

31. Mayer, A., Scheglmann, D., Dove, S., Glatz, A., Wickner, W., and Haas, A. (2000). Phosphatidylinositol 4,5-bisphosphate regulates two steps of homotypic vacuole fusion. Mol Biol Cell 11, 807–17.

32. Mayer, A., Wickner, W., and Haas, A. (1996). Sec18p (NSF)-driven release of Sec17p (alpha-SNAP) can precede docking and fusion of yeast vacuoles. Cell 85, 83–94.

33. Miner, G. E., Starr, M. L., Hurst, L. R., and Fratti, R. A. (2017). Deleting the DAG kinase Dgk1 augments yeast vacuole fusion through increased Ypt7 activity and altered membrane fluidity. Traffic 18, 315–329.

34. Miner, G. E., Starr, M. L., Hurst, L. R., Sparks, R. P., Padolina, M., and Fratti, R. A. (2016). The Central Polybasic Region of the Soluble SNARE (Soluble N-Ethylmaleimide-sensitive Factor Attachment Protein Receptor) Vam7 Affects Binding to Phosphatidylinositol 3-Phosphate by the PX (Phox Homology) Domain. J Biol Chem 291, 17651–17663.

35. Miyamoto, S., and Kollman, P. A. (1992). SETTLE: An Analytical Version of the SHAKE and RATTLE Algorithm for Rigid Water Molecules. J Comput Chem 13, 952–962.

36. Nakanishi, H., Morishita, M., Schwartz, C. L., Coluccio, A., Engebrecht, J., and Neiman, A. M. (2006). Phospholipase D and the SNARE Sso1p are necessary for vesicle fusion during sporulation in yeast. J Cell Sci 119, 1406–1415.

37. Pande, V. S., Beauchamp, K., and Bowman, G. R. (2010). Everything you wanted to know about Markov State Models but were afraid to ask. Methods 52, 99–105.

38. Phillips, J. C., Braun, R., Wang, W., Gumbart, J., Tajkhorshid, E., Villa, E., Chipot, C., Skeel, R. D., Kalé, L., and Schulten, K. (2005). Scalable molecular dynamics with NAMD. J Comput Chem 26, 1781–1802.

39. Putta, P., Rankenberg, J., Korver, R. A., van Wijk, R., Munnik, T., Testerink, C., and Kooijman, E. E. (2016). Phosphatidic acid binding proteins display differential binding as a function of membrane curvature stress and chemical properties. Biochim Biophys Acta 1858, 2709–2716.

40. Roberts, R. L., Barbieri, M. A., Pryse, K. M., Chua, M., Morisaki, J. H., and Stahl, P. D. (1999). Endosome fusion in living cells overexpressing GFP-rab5. J Cell Sci 112, 3667–3675.

41. Rogasevskaia, T. P., and Coorssen, J. R. (2015). The Role of Phospholipase D in Regulated Exocytosis. J Biol Chem 290, 28683–28696.

42. Ryu, J. K., Min, D., Rah, S. H., Kim, S. J., Park, Y., Kim, H., Hyeon, C., Kim, H. M., Jahn, R., and Yoon, T. Y. (2015). Spring-loaded unraveling of a single SNARE complex by NSF in one round of ATP turnover. Science 347, 1485–1489.

43. Sali, A., and Blundell, T. L. (1993). Comparative protein modelling by satisfaction of spatial restraints. J Mol Biol 234, 779–815.

44. Sasser, T., Qiu, Q. S., Karunakaran, S., Padolina, M., Reyes, A., Flood, B., Smith, S., Gonzales, C., and Fratti, R. A. (2012). Yeast lipin 1 orthologue pah1p regulates vacuole homeostasis and membrane fusion. J Biol Chem 287, 2221–2236.

45. Schreiner, E., Trabuco, L. G., Freddolino, P. L., and Schulten, K. (2011). Stereochemical errors and their implications for molecular dynamics simulations. BMC Bioinformatics 12, 190.

46. Sollner, T., Bennett, M. K., Whiteheart, S. W., Scheller, R. H., and Rothman, J. E. (1993). A protein assembly-disassembly pathway in vitro that may correspond to sequential steps of synaptic vesicle docking, activation, and fusion. Cell 75, 409–418.

47. Sparks, R. P., Jenkins, J. L., Miner, G. E., Wang, Y., Guida, W. C., Sparks, C. E., Fratti, R. A., and Sparks, J. D. (2016). Phosphatidylinositol (3,4,5)-trisphosphate binds to sortilin and competes with neurotensin: Implications for very low density lipoprotein binding. Biochem Biophys Res Commun 479, 551–556.

48. Starr, M. L., Hurst, L. R., and Fratti, R. A. (2016). Phosphatidic acid sequesters Sec18p from cis-SNARE complexes to inhibit priming. Traffic 17, 1091–1109.

49. Stroupe, C., Collins, K. M., Fratti, R. A., and Wickner, W. (2006). Purification of active HOPS complex reveals its affinities for phosphoinositides and the SNARE Vam7p. Embo J 25, 1579–1589.

50. Studier, F. W. (2014). Stable expression clones and auto-induction for protein production in E. coli. Methods Mol Biol 1091, 17–32.

51. Trabuco, L. G., Villa, E., Mitra, K., Frank, J., and Schulten, K. (2008). Flexible fitting of atomic structures into electron microscopy maps using molecular dynamics. Structure 16, 673–683.

52. Tudyka, T., and Skerra, A. (1997). Glutathione S-transferase can be used as a C-terminal, enzymatically active dimerization module for a recombinant protease inhibitor, and functionally secreted into the periplasm of Escherichia coli. Protein Sci 6, 2180–2187.

53. Vollrath, F., Hawkins, N., Porter, D., Holland, C., and Boulet-Audet, M. (2014). Differential Scanning Fluorimetry provides high throughput data on silk protein transitions. Sci Rep 4, 5625.

54. Wilson, D. W., Whiteheart, S. W., Wiedmann, M., Brunner, M., and Rothman, J. E. (1992). A multisubunit particle implicated in membrane fusion. J Cell Biol 117, 531–538.

55. Yavuz, H., Kattan, I., Hernandez, J. M., Hofnagel, O., Witkowska, A., Raunser, S., Walla, P. J., and Jahn, R. (2018). Arrest of *trans-SNARE* zippering uncovers loosely and tightly docked intermediates in membrane fusion. J Biol Chem 293, 8645–8655.

56. Yu, R. C., Hanson, P. I., Jahn, R., and Brünger, A. T. (1998). Structure of the ATP-dependent oligomerization domain of N-ethylmaleimide sensitive factor complexed with ATP. Nat Struct Biol 5, 803–811.

57. Yu, R. C., Jahn, R., and Brunger, A. T. (1999). NSF N-terminal domain crystal structure: models of NSF function. Mol Cell 4, 97–107.

58. Zhao, M., and Brunger, A. T. (2016). Recent Advances in Deciphering the Structure and Molecular Mechanism of the AAA+ ATPase N-Ethylmaleimide-Sensitive Factor (NSF). J Mol Biol 428, 1912–1926.

59. Zhao, M., Wu, S., Zhou, Q., Vivona, S., Cipriano, D. J., Cheng, Y., and Brunger, A. T. (2015). Mechanistic insights into the recycling machine of the SNARE complex. Nature 518, 61–67.

